# DNA uptake and twitching motility are controlled by the small RNA Arp through repression of pilin translation in *Acinetobacter baumannii*

**DOI:** 10.1101/2025.07.19.665661

**Authors:** Fergal J. Hamrock, Thomas Guest, Mikaela N. Daum, Orla Connell, Anna S. Ershova, Karsten Hokamp, Alastair B. Fleming, Michael J. Gebhardt, Alexander J. Westermann, Carsten Kröger

## Abstract

*Acinetobacter baumannii* is a major opportunistic pathogen capable of natural transformation, a process driven by type IV pili (T4P) that facilitates horizontal gene transfer and accelerates the spread of antimicrobial resistance. While the transcriptional regulation of T4P is increasingly understood, post-transcriptional mechanisms controlling pilus assembly remain unexplored. Here, we identify and characterise a small RNA, Arp (*Acinetobacter* repressor of pilin), as a post-transcriptional repressor of T4P-mediated functions in *A. baumannii*. In a previous Hi-GRIL-seq experiment, we detected specific ligation events between Arp and the ribosome binding site of the *pilA* mRNA, encoding the major pilin subunit PilA. In-line probing and translational reporter assays revealed that Arp represses *pilA* translation by sequestering the Shine-Dalgarno sequence and the first 17 codons of the mRNA. Overexpression of Arp significantly impairs DNA uptake and twitching motility, two hallmark T4P-dependent phenotypes. Together, our findings identify a native *A. baumannii* sRNA that modulates natural competence by targeting pilin synthesis, revealing a new regulatory layer that could be exploited to disrupt horizontal gene transfer in multidrug-resistant strains.

**Significance Statement:** *Acinetobacter baumannii* is a multidrug-resistant WHO #1 priority pathogen that acquires antibiotic resistance genes through natural transformation, a process dependent on type IV pili (T4P). This work reveals Arp, the first native post-transcriptional repressor of natural competence in *A. baumannii*, uncovering a novel regulatory layer that modulates horizontal gene transfer. The widespread presence of *arp* in pathogenic *Acinetobacter* strains suggests that sRNA is an important regulator in those organisms. Furthermore, these findings broaden our understanding of RNA-based regulation in this priority pathogen and open potential avenues for interfering with antibiotic resistance dissemination.

## Introduction

*Acinetobacter baumannii* is an opportunistic pathogen that primarily affects critically ill, hospitalised patients (1). It is associated with high-mortality infections such as ventilator-associated pneumonia, bloodstream infections, and catheter-related urinary tract infections (2, 3). Infections are often complicated by extensive antimicrobial resistance (AMR), enabling *A. baumannii* to persist in clinical settings and evade chemotherapeutic treatment; accordingly, public health agencies have designated carbapenem-resistant *A. baumannii* a priority pathogen in urgent need of novel therapeutic strategies (4). Horizontal gene transfer (HGT) is a major contributor to AMR in clinical settings and allows many bacterial pathogens, including *A. baumannii*, to acquire antimicrobial resistance genes (ARGs) and virulence factors (5–7). While conjugation is a key HGT mechanism, natural transformation also plays a central role in the dissemination of resistance determinants and genomic diversification (8–11). Many clinical strains of *A. baumannii* exhibit natural competence, enabling them to internalise exogenous DNA, including both plasmids and linear fragments (12). Exogenous DNA can also be integrated into the chromosome, provided sufficient sequence homology, through homologous recombination (13–15).

Natural competence in *A. baumannii* is primarily mediated by the type IV pilus (T4P), a dynamic surface appendage essential for DNA uptake across the outer membrane. The pili also facilitate twitching motility on solid surfaces, linking surface movement with the ability to acquire DNA from the environment (16–18). T4P are composed of multiple protein subunits, including PilA — the major pilin protein in *Acinetobacter spp.* which displays sequence and structural diversity (19). While the regulation of T4P is an active area of investigation, transformation frequencies have been shown to be sensitive to specific environmental conditions including growth phase (18), calcium ions, serum albumin, subinhibitory concentrations of antibiotics and the PilSR two-component system (18, 20–22). Despite these advances in understanding transcriptional control of T4P, the complexity of regulation is incompletely understood in *A. baumannii*.

Bacterial small RNAs (sRNAs) are key post-transcriptional regulators. While HGT is considered a major contributor to sRNA evolution and spread (23–25), the activity of certain sRNAs was in turn found to impact natural competence in several bacterial species (26–28). Mechanistically, sRNAs can modulate gene expression by forming imperfect base-pairing interactions with target transcripts, typically within the 5′ untranslated region (5′ UTR) of messenger RNAs (29, 30). These base-pairing events are frequently initiated by sRNA “seed regions” that bind to complementary nucleotides within the target mRNA (31). These sRNA-mRNA interactions often result in repression of translation by occluding the Shine-Dalgarno (SD) sequence and adjacent coding region, thereby blocking ribosome access or by accelerating degradation of the target mRNA (29, 30).

Using Hi-GRIL-seq, a proximity ligation method to identify sRNA-mRNA interactions in *A. baumannii* AB5075 independently of the RNA binding proteins, we previously identified numerous potential sRNA-target pairs (32, 33). One sRNA candidate, designated sRNA40 and hereafter referred to as *Acinetobacter* repressor of pilin (Arp), was repeatedly ligated to the translational initiation site of the *pilA* mRNA. The existence of chimeric Arp-*pilA* mRNA molecules suggested that Arp may engage in base-pairing with the ribosome binding site (RBS) of *pilA*, potentially interfering with translation and impacting T4P-mediated phenotypes. In this study, we investigated the role of Arp in controlling DNA uptake and twitching motility, which are key mechanisms of horizontal gene transfer in *A. baumannii*.

## Results

### Hi-GRIL-seq identifies Arp-*pilA* mRNA interaction in *A. baumannii* AB5075

Previously, we used Hi-GRIL-seq of *A. baumannii* AB5075 in three different growth conditions and identified hundreds of potential sRNA-mRNA interactions that might represent events of sRNA-mediated post-transcriptional regulation of translation (33). In this experiment, 119 RNA chimeras were detected where the sRNA candidate sRNA40 (Arp) was ligated to *pilA* mRNA, which encodes the pilin protein of *A. baumannii*. Arp-*pilA* chimeras were more numerous than the two other potential interaction partners that were detected in the Hi-GRIL-seq experiment (namely with *ABUW_RS10585*, encoding an acetyltransferase and *ABUW_RS18070*, the mRNA of a tRNA-Ala molecule (**Supplementary** Figure 1A)). However, unlike the interaction with *pilA*, these ligation events had low abundance, with only 11 and 10 chimeric molecules, respectively. To facilitate the visualization of Arp chimeras in the Hi-GRIL-seq data, we provide an interactive online browser, accessible at: http://bioinf.gen.tcd.ie/jbrowse2/Hi-GRIL-seq/Arp. Bioinformatic prediction suggested the formation of a long high affinity RNA-RNA duplex (hybridisation energy= -50.18 kcal/mol) between Arp and *pilA* mRNA extensively covering the SD and the first seven codons of *pilA* (**Figure 1A**). In our previous Hi-GRIL-seq study, chimeras were detected under four distinct conditions: a non-induced (NI) and an induced (IND) T4 RNA ligase control, as well as following T4 RNA ligase induction after iron starvation with 2,2′-dipyridyl (DIP) or antibiotic stress with imipenem (IMIP) (33). More chimeras were observed in the IND (n=40), DIP (n=50) and IMIP (n=28) conditions of which 88.24% of these chimeric reads mapped to the predicted interaction site (n=105), while the T4 RNA ligase non-induced control experiment only contained one chimera **(Figure 1B)**.

**Figure 1.**
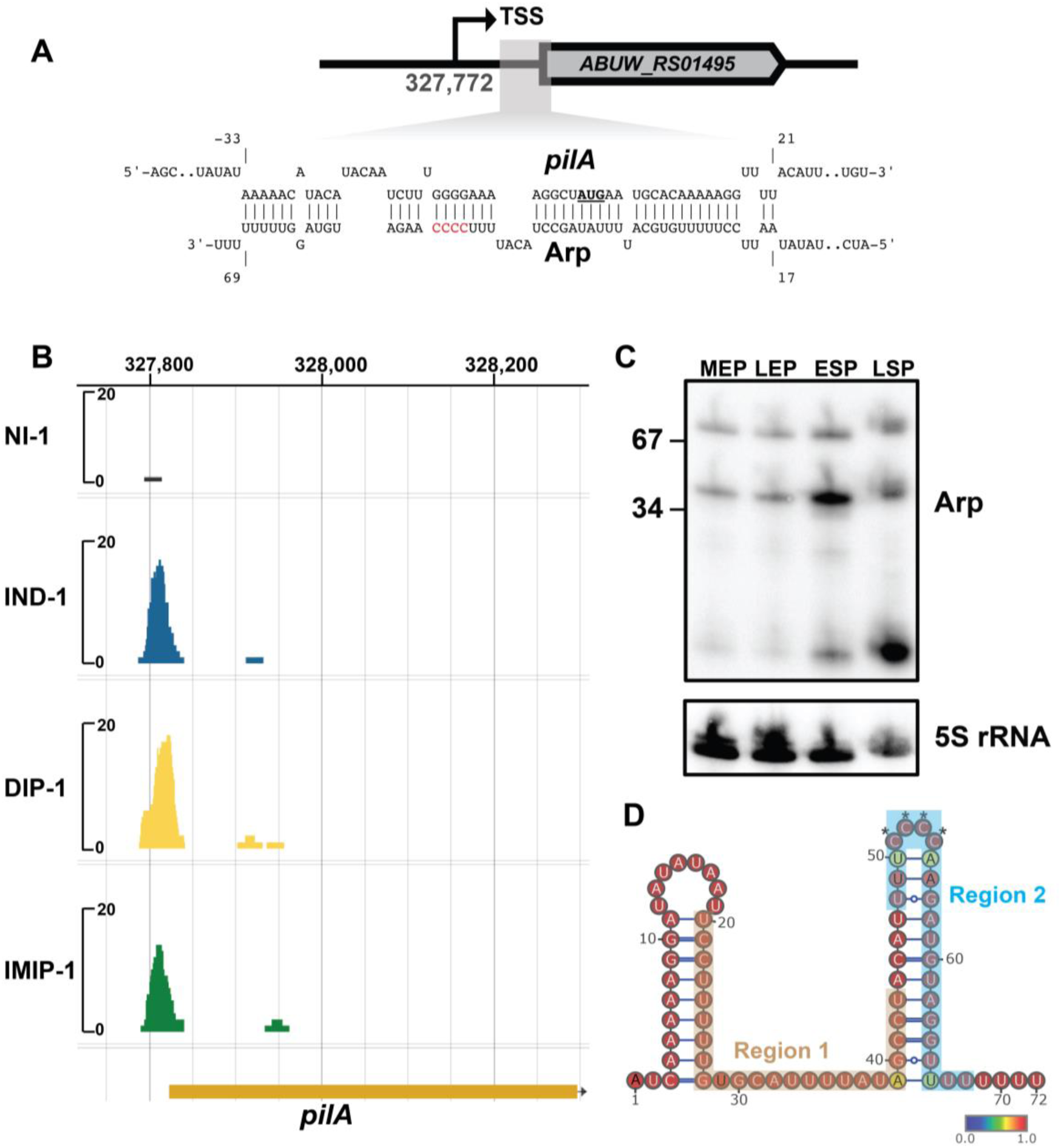
Identification of Arp sRNA-*pilA* mRNA interactions in *A. baumannii*. (**A**) The sRNA-mRNA interaction prediction tool IntaRNA (75) was used to identify the genomic location of *Arp*-*pilA* mRNA base-pairing interactions. The genomic location of the predicted interaction duplex is shown relative to *ABUW_RS01495* (*pilA*). The positions of *pilA* nucleotides are numbered relative to the translational initiation site, where the *pilA* start codon (AUG) is emboldened and underlined. The potential Arp interacting region is highlighted in red font. The transcriptional start site (TSS) of *ABUW_RS01495* (*pilA*) is also indicated. (**B**) The locations of the *pilA* portions of Arp-*pilA* chimeras that were detected by Hi-GRIL-seq were mapped to the AB5075 chromosome for each condition used in that study. This includes the T4 RNA ligase non-induced control (NI), induced T4 RNA ligase control (IND), 2′,2-dipyridyl iron starvation condition (DIP) and the imipenem shock condition (IMIP). (**C**) The expression of Arp across *A. baumannii* AB5075 growth was assessed by northern blotting with RNA isolated from mid exponential phase (MEP), late exponential phase (LEP), early stationary phase (ESP) and late stationary phase (LSP). (**D**) The minimum free energy (MFE) structure of Arp was predicted using RNAfold and was visualised using VARNA (76, 77). All nucleotides are colour based on their base-pairing probabilities. The nucleotides involved in base-pairing (Regions 1 and 2) are labelled). The Arp interacting region is highlighted with asterisks (*).

Differential RNA-sequencing (dRNA-seq) in *A. baumannii* ATCC17978 previously determined that *arp* encodes both a ∼96-nucleotide and ∼72-nucleotide long sRNA (34). However, Arp appears truncated at the 5′ end in the Hi-GRIL-seq RNA-seq reads in *A. baumannii* AB5075, indicating that it forms the shorter sRNA isoform (**Supplementary** Figure 1B). To examine the size and expression profile of this sRNA in *A. baumannii* AB5075, northern blotting to detect Arp expression levels and any other isoforms was done with RNA isolated from various stages of growth in rich medium (L-broth) including mid exponential phase (MEP, OD_600_=0.3), late exponential phase (LEP, OD_600_=1), early stationary phase (ESP, OD_600_=2) and late stationary phase (LSP, 16 h of growth). This revealed that Arp transcripts accumulate throughout growth, attaining the highest levels of abundance in ESP and LSP (**Figure 1C**). The stable ∼72 nt long full-length RNA in the AB5075 strain aligns with the Hi-GRIL-seq findings. In addition to 72 nt Arp, smaller isoforms accumulate in ESP and LSP (the Hi-GRIL-seq experiment had been carried out at ESP). It is unclear whether these are functional or products of degradation, but it might suggest that Arp exerts its regulatory impact in the later stages of growth. As inferred from RNA-seq read coverage profiles, there were no obvious processing sites within Arp (**Supplementary** Figure 1B). *In-silico* prediction of the 72 nt Arp secondary structure suggests it forms two hairpins, one at the 5′ end and the other at the 3′ end flanked by a poly(U) tail to promote Rho-independent termination (**Figure 1D**). Approximately 61% of nucleotides (n=44) are predicted to be involved in the Arp-*pilA* duplex structure (**Figure 1A**), suggesting that unfolding of both hairpin loops of the sRNA would be needed for interaction with *pilA* mRNA. We predicted that the Arp interaction region is located at the apex of the 3′ terminator stem-loop structure because of its single-strandedness that could base-pair with another RNA molecule as a potential seed region, an uncommon—yet not unprecedented (35, 36)—feature within bacterial sRNAs.

### Arp is a conserved sRNA among *Acinetobacter* species

To investigate the conservation of Arp and its regulatory architecture among *Acinetobacter* species, we aligned sequences of *arp* and its upstream region (128 nt) from nine representative *A. baumannii* strains and *A. nosocomialis* (**Supplementary** Figure 2A). We observed a high degree of sequence conservation, particularly within the sRNA gene. While *A. nosocomialis* displayed the most divergence in the upstream sequence, the core sRNA structure remained mostly intact. Notably, no clear homologue of *arp* was found in *A. baylyi*, a non-pathogenic relative. Putative promoter elements were identified upstream of *arp* in multiple strains. Notably, a ChIP-seq study in *A. baumannii* ATCC17978 identified a BfmR binding motif located upstream of *arp* and RNA-seq in *A. baumannii* ATCC17961 showed that *arp* expression was down-regulated in stationary phase in a Δ*bfmR* mutant indicating that BfmR could be a direct activator of the expression of *arp* (37, 38). This region is conserved among the strains used in this analysis and is depicted (**Supplementary** Figure 2A).

To expand this analysis, we used a BLAST-based approach to search for *arp* homologues in 923 available, complete *Acinetobacter* genomes. This broader screen provided a comprehensive view of *arp* distribution across the genus (**Supplementary** Figure 2B). We identified 725 hits with >95% identity and >90% query coverage, indicating that *arp* is highly conserved across a substantial portion of the genus. Notably, while *arp* is not universally present in *A. baumannii*, it is completely identical in 551 genomes, suggesting strong conservation in a specific subset of strains. This apparent ’all-or-nothing’ pattern may reflect a lineage-specific adaptation potentially involved in modulating twitching motility and DNA uptake, or alternatively, a relatively recent emergence of the sRNA that is spreading through HGT.

### Arp interacts with *pilA in vitro* using a contiguous interacting region

To determine whether Arp interacts with *pilA*, electrophoretic mobility shift assays (EMSAs) were conducted using *in vitro* transcribed and radiolabelled Arp with increasing concentrations of a 102-nt *pilA* RNA fragment (**Figure 2**). The Arp RNA fragment began at the native transcriptional start site (TSS) and included the predicted Arp interaction site. As *pilA* concentration increased, a clear band shift was observed, indicating strong base-pairing interactions *in vitro* (*K*_D_ = 29.02 nM; **Figure 2A**, **Figure 2D**). To assess the role of the predicted Arp interacting region in *pilA* binding, the region was mutated from 5′-CCCC-3′ to 5′-GGGG-3′, generating Arp* (**Figure 1D**). EMSAs with Arp* and *pilA* showed a strong reduction in band shifting (*K*_D_ = 139.9 nM; **Figure 2B**, **Figure 2D**), suggesting that whilst the potential interacting region contributes to binding, additional nucleotides are also involved, consistent with the extended interaction predicted in **Figure 1A**. When Arp* was incubated with a compensatory *pilA* mutant (*pilA*\*, mutated from 5′-GGGG-3′ to 5′-CCCC-3′), high-affinity binding was restored (*K*_D_ = 7.89 nM; **Figure 2C**, **Figure 2D**), supporting the conclusion that the interaction is sequence-specific, driven by base-pairing and highly dependent on the four-GC duplex.

**Figure 2.**
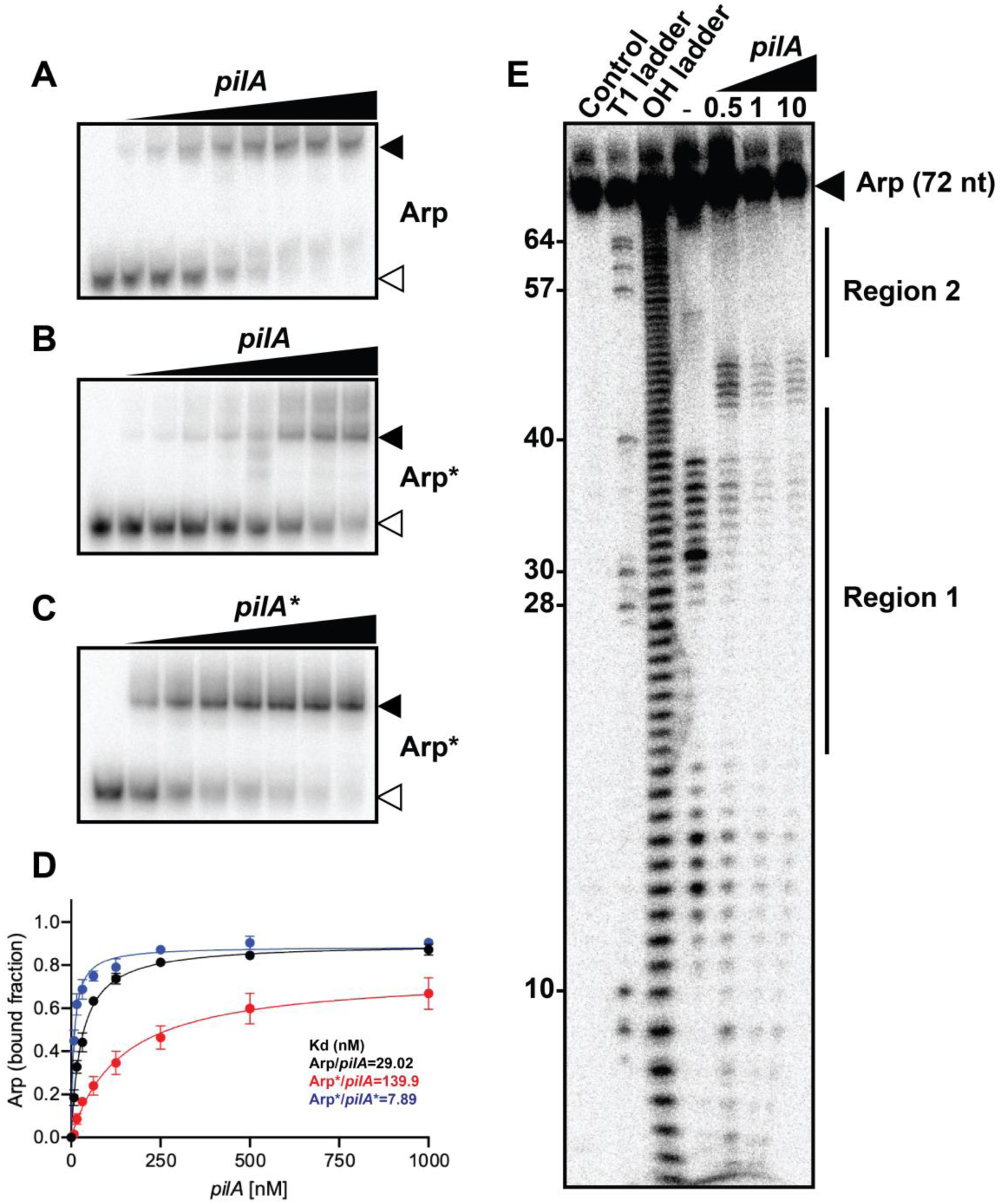
Arp interacts with *pilA* RNA *in vitro*. (**A**) The ability of 5′-radiolabelled wildtype Arp and (**B**) interacting region mutated Arp (Arp*) to form base-pairing interactions with increasing concentrations of *pilA* or (**C**) *pilA\** was assessed using electrophoretic motility shift assays. This resulted in clear band shifts (black arrow) that were distinct from the full-length sRNA (white arrow). (**D**) EMSAs were used to quantify the dissociation constants *K_D_* of these targets. The *K_D_* values were calculated from the mean of four independent replicates (n=4). (**E**) The exact Arp nucleotides involved in *pilA* base-pairing were identified using in-line probing. The protected regions (Region 1 and Region 2) are indicated.

To determine the Arp secondary structure experimentally and identify nucleotides involved in *pilA* base-pairing, in-line probing was performed. When Arp was incubated in the absence of *pilA*, the sRNA adopted a secondary structure matching the Arp *in silico* prediction that consists of two stem-loop structures at the 5′ and 3′-ends linked by a short, single stranded region (**Figure 1D**, **Figure 2E**). In-line probing also revealed that two adjacent regions of Arp; Region 1 and Region 2 (the Arp interacting region is encompassed within Region 2), and corresponding to nucleotides 20-43 and 48-68, were involved in *pilA* base-pairing (**Figure 2E)**. The RNA duplex structure determined *in vitro* therefore matches the extensive base-pairing prediction (**Figure 1A**).

### Arp represses PilA translation in a heterologous reporter model

To determine functional consequences of Arp-*pilA* interactions on PilA expression, we used an established translational reporter system in *E. coli* (39, 40). This heterologous model has previously been used to validate sRNA-mRNA interactions from other species, including *A. baumannii* (33, 41). By measuring the relative fluorescence of *E. coli* constitutively expressing PilA-sfGFP translational fusions, in the presence or absence of Arp, the impact of Arp on the translation of PilA-sfGFP can be inferred. Expression of Arp significantly reduced the fluorescence intensity of PilA-sfGFP (∼10-fold repression, p<0.0001), suggesting that Arp represses the translation of PilA *in vivo* (**Figure 3A**). The Arp* mutant partially disrupted repression of PilA-sfGFP fluorescence intensity (∼2.6-fold repression, p<0.01) compared to a strain overexpressing wild-type Arp, indicating that additional nucleotides are involved in Arp base-pairing outside of this potential interacting region, in agreement with the EMSAs (**Figure 2**). The disruption of the predicted Arp binding site in the *pilA*\* compensatory mutants rendered the translational fusion constructs non-functional due to disruption of the *pilA* SD sequence (**Figure 3A**).

**Figure 3.**
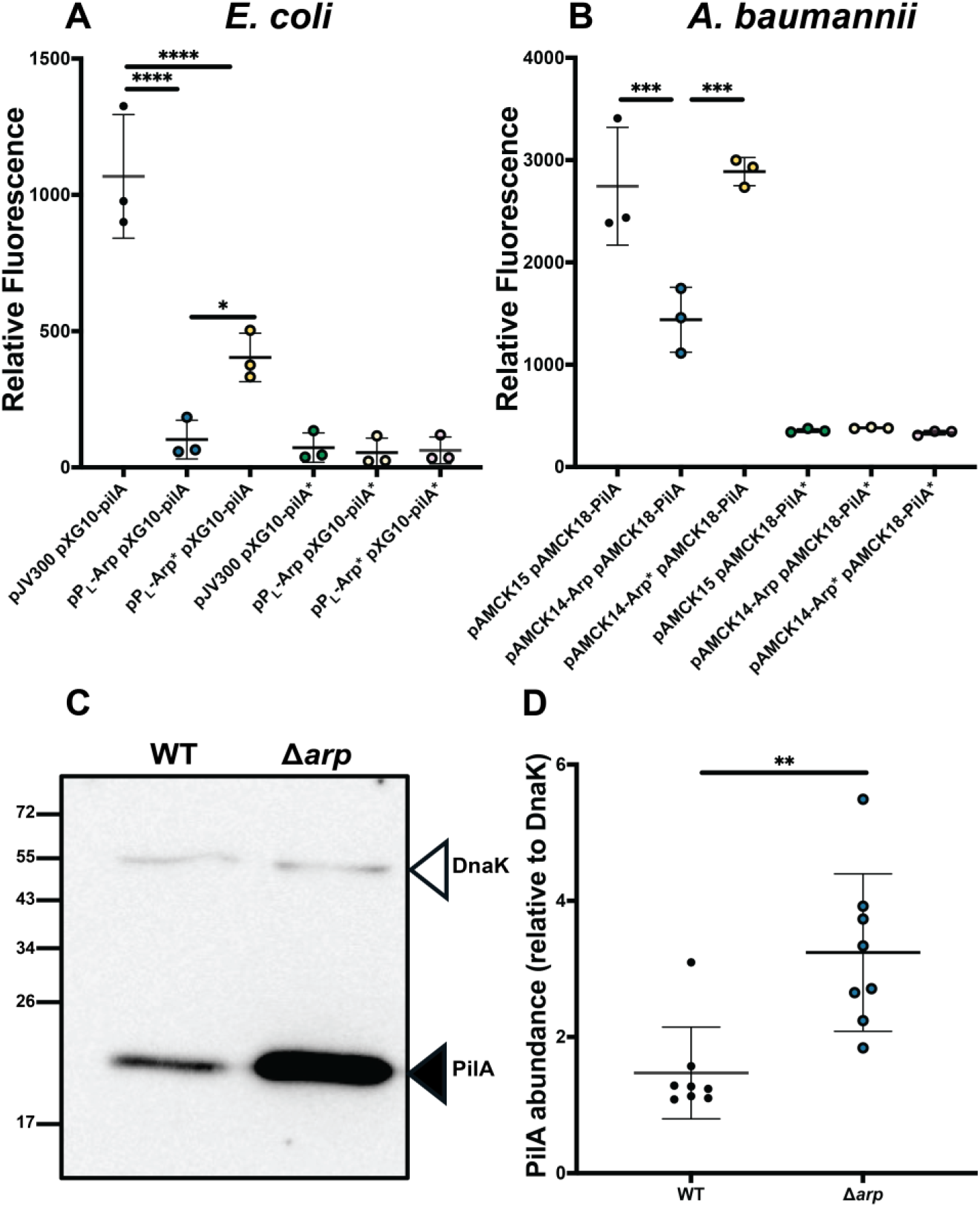
Arp inhibits PilA-sfGFP translation. Overexpression of Arp reduces PilA-sfGFP expression *in vivo* in both (**A**) *Escherichia coli* and (**B**) *Acinetobacter baumannii* AB5075, as measured by PilA-sfGFP translational reporters. (**C**) Endogenous Arp similarly suppresses PilA-FLAG steady-state levels during growth on solid media, with higher PilA-FLAG abundance observed in the Δ*arp* mutant relative to the wildtype (WT). (**D**) Quantification of PilA-FLAG levels normalised to the DnaK loading control. Error bars represent the standard deviation of three independent biological replicates (n=3) for translational reporter assays and eight biological replicates (n=8) for western blot quantification. Statistical analysis was performed using one-way ANOVA with Tukey’s post hoc test for translational reporter data, and an unpaired t-test for western blot comparisons. Significance was defined where * denotes *P*<0.05, ** denotes *P*<0.01, *** denotes *P*<0.001 and **** denotes *P*<0.0001.

### Arp inhibits translation of PilA *in vivo* in *A. baumannii*

To assess whether Arp functions as a post-transcriptional repressor *in vivo*, an alternative translational reporter system was used which has been validated for confirming sRNA-mRNA interactions in *A. baumannii* (42). Like the pXG10sf plasmid used in *E. coli* (**Figure 3A**), expression of a PilA-sfGFP translational fusion is driven by the constitutive P_LtetO_ promoter on plasmid pAMCK18-PilA. This reporter strain was transformed with either a control plasmid (pAMCK15) or an Arp-overexpression construct (pAMCK14-Arp), in which Arp is constitutively expressed from the P_LlacO_ promoter (the same promoter as in pP_L_ plasmid used in *E. coli*). Consistent with the data in *E. coli*, Arp overexpression significantly reduced PilA-sfGFP fluorescence (∼1.9-fold, p < 0.001) compared to the control strain in Δ*arp A. baumannii* and repression was fully relieved by mutation of the Arp potential interacting region (Arp*), which restored fluorescence to wild-type levels (**Figure 3B**). The essential nature of Hfq in *A. baumannii* AB5075 complicates efforts to determine its role in facilitating sRNA-mRNA base-pairing *in vivo* using this reporter system (43, 44). To assess whether Hfq contributes to these interactions, we searched for ARN motifs—known Hfq-binding sites—within the Arp sequence. We identified 10 such motifs, with half located within the 5′ stem-loop structure adjacent to Region 1 of Arp (**Supplementary** Figure 3), suggesting that Hfq may indeed participate in mediating base-pairing. To determine whether endogenous Arp expression is sufficient to modulate PilA protein levels *in vivo*, PilA levels were determined using *A. baumannii* strains carrying a chromosomal PilA::3×FLAG C-terminal fusion in both wildtype and Δ*arp* backgrounds by western blotting (**Supplementary** Figure 4). The strains were grown in motility medium, known to promote twitching motility by inducing expression of PilA on semi-solid media (16). A significant increase in PilA::3×FLAG levels in the Δ*arp* strain was observed relative to wildtype (2.2-fold, p=0.0022; **Figure 3C** and **D**), demonstrating that native levels of Arp are sufficient to repress PilA protein accumulation under physiological conditions. The result strongly supports a model in which Arp inhibits translation of PilA by blocking ribosome access to the initiation region. Taken together, the data collected using both artificial and native expression systems, indicate that Arp represses PilA production at the level of translation *in vivo*, which ultimately results in measurable consequences on PilA protein production.

### Arp restricts motility in *A. baumannii*

As PilA is an essential component of the T4P in *A. baumannii*, crucial for movement on solid surfaces, the role of Arp in modulating twitching motility was assessed. We compared wild-type and Δ*arp A. baumannii* strains carrying the pAMCK15 control vector along with the non-motile PilA deletion mutant (Δ*pilA*), an Arp-overexpression strain (Δ*arp* carrying pAMCK14-Arp), and an Arp-overexpression strain with the interacting region mutated (Δ*arp* carrying pAMCK14-Arp*). To determine whether Arp represses twitching motility, individual colonies were inoculated at the centre of motility plates, incubated upright at 37°C for five days and the resulting motility zones were measured (**Figure 4A and 4B**). The Δ*arp* strain exhibited a ∼2-fold increase in motility relative to wild-type (p<0.0001), suggesting that Arp reduces motility.

**Figure 4.**
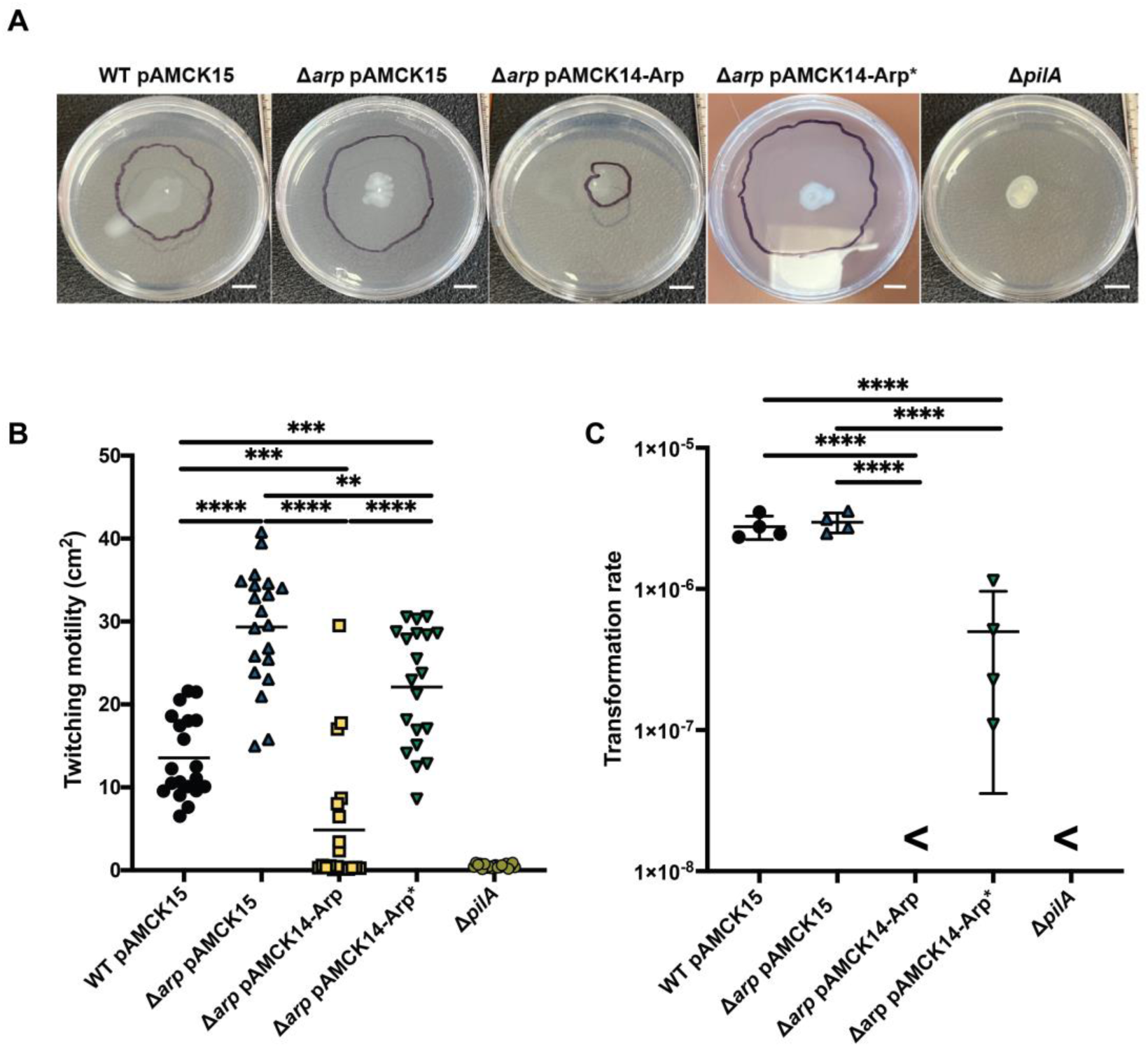
Arp regulates twitching motility and natural transformation in *A. baumannii.* (**A**) Representative images of twitch motility plates and (**B**) quantification of twitching motility zones in *A. baumannii* AB5075 wild-type (WT) and Arp-deletion strains (Δ*arp*) carrying either an empty vector (pAMCK15), an Arp overexpression plasmid (pAMCK14-Arp), or a interacting region mutant of Arp (pAMCK14-Arp*). A Δ*pilA* strain was included as a negative control. For each of the twitching motility plates, the motility zones are outlined; scale bars indicate 1 cm. (**C**) Natural transformation frequencies of the same strains assessed by uptake of the pAMCK18-PilA plasmid. No transformants were recovered on tetracycline-containing L-plates for the Arp-overexpression strain or the Δ*pilA* control (denoted by <). Error bars represent the standard deviation of twenty independent biological replicates (n=20) for the twitching motility assay and four independent biological replicates (n=4) for the natural transformation assay. Statistical analysis was performed using one-way ANOVA with Tukey’s post hoc test. Significance was defined where * denotes *P*<0.05, ** denotes *P*<0.01, *** denotes *P*<0.001 and **** denotes *P*<0.0001.

Conversely, Arp overexpression significantly reduced motility compared to wild-type and Δ*arp* (∼3-fold and ∼6-fold reductions, respectively; p<0.0001), almost mirroring the non-motile phenotype of the Δ*pilA* mutant. The disruption of the Arp interacting region in this construct significantly increased twitching motility (∼5-fold, p<0.0001) highlighting the importance of an intact interacting region in initiating base-pairing with *pilA* mRNA. Together, these results demonstrate that Arp is a physiologically relevant post-transcriptional repressor of twitching motility in *A. baumannii*.

### Arp inhibits DNA uptake in *A. baumannii* AB5075

Given the importance of T4P in enabling natural transformation in *A. baumannii*, Arp may also impact on DNA uptake. The same panel of strains used in the twitching motility assay were thus used to assess transformation efficiency following overnight incubations with plasmid DNA (pAMCK18-PilA) in transformation medium. Transformation frequencies were determined by plating the resulting cells on selective and non-selective L-agar (**Figure 4C**). There were comparable colony counts between strains on the non-selective plates, confirming consistent growth (**Supplementary** Figure 5). No transformants were recovered from the Arp overexpression strain, indicating a significant reduction of DNA uptake (p<0.0001) compared to the wild-type and Δ*arp* strains. In the Δ*arp* Arp* overexpression strain, DNA uptake was partially restored, indicating that residual binding capacity of Arp* with *pilA* mRNA impacts DNA uptake but is still demonstrably higher compared to overexpression of wild-type Arp. These findings illustrate that high Arp expression is sufficient to inhibit natural transformation in *A. baumannii*, likely due to post-transcriptional repression of PilA, and thus, the DNA uptake machinery in this pathogen.

## Discussion

The emergence of multidrug-resistant *A. baumannii* strains is driven by HGT, a process tightly linked to T4P-mediated DNA uptake by natural transformation and twitching motility (16–18). In this study, Arp, a previously uncharacterised small RNA, was discovered as a post-transcriptional regulator of *pilA*, the major pilin subunit of the T4P machinery. Arp represses *pilA* translation through direct base-pairing, thereby reducing twitching motility and DNA uptake. The characterisation of Arp uncovers a layer of post-transcriptional control of HGT, which is highly relevant to AMR evolution, because DNA uptake and twitching motility are tightly interconnected (17, 45, 46). Equally, delineating the mechanistic details of repression of PilA translation by Arp expands the understanding of sRNA-mediated post-transcriptional regulation in *A. baumannii*, which is still poorly defined (33, 47). Only recently, the mechanistic understanding of small RNA-mediated gene regulation in *A. baumannii* has deepened by uncovering the regulation of *carO* and *bfnH* by the sRNA Aar (33) and type-VI secretion system gene *tssM* by sRNA AbsR28 (48). Although the proximity ligation-based Hi-GRIL-seq is a relatively crude tool to identify sRNA-mRNA interactions, it identified numerous Arp-*pilA* chimeras that reflect *bona fide* post-transcriptional regulatory events *in vivo*. These base-pairing events were localised to the translational initiation site of the *pilA* transcript, indicating that Arp acts as a canonical repressor of PilA translation, much like the mechanism of Aar in *A. baumannii* and most of the characterized sRNAs in other bacteria (30, 33). The computational prediction and *in vitro* interaction of *pilA* mRNA (**Figures 1 and 2**) and Arp proved to hold true *in vivo* (**Figures 3 and 4**), and we identified the potential Arp interacting region consisting of a stretch of four cytosine nucleotides located in the tip of the second loop critical for interactions with *pilA*. Importantly, native levels of Arp were sufficient to impair *A. baumannii* twitching motility and overexpression of Arp suppresses this phenotype along with DNA uptake in a base-pairing-dependent manner (**Figure 4**). Arp may represent an atypical mode of sRNA-mediated repression, as the structured interacting region overlaps with the terminal stem-loop rather than lying in a linear region, which is reminiscent of the mechanism of ManS where a seed region is part of the head of a terminator loop (36). In summary, we establish Arp as the first post-transcriptional repressor of natural competence in *A. baumannii*.

Arp is the latest member of a group of sRNAs that regulate natural transformation among a handful of bacterial species, including *Legionella pneumophila*, *Vibrio cholerae* and *Streptococcus pneumoniae* (26–28). Arp bears strongest conceptual similarity with *L. pneumophila* sRNA RocR, that represses natural transformation by binding an mRNA critical for DNA uptake, the *comEA* 5′-UTR (28). Like RocR, Arp directly represses DNA uptake machinery, rather than regulating upstream activators of the competence pathway. However, unlike RocR, Arp forms more extensive base-pairing interactions that occlude the entire translational initiation region of its target transcript. While RocR requires the FinO family RNA chaperone RocC to exert its repressive activity in *L. pneumophila* (28, 49), the role of any chaperone in facilitating Arp function remains unclear. While our EMSA and in-line probing data (**Figure 3**) suggest that Arp can bind *pilA* mRNA independently *in vitro*, this does not rule out the involvement of an RNA-binding protein such as Hfq *in vivo*. Recently, Hfq was suggested to be an essential gene in *A. baumannii* AB5075 (used in this study) as efforts to delete *hfq* repeatedly failed (43), and this was supported by a later CRISPR-interference study showing that the knockdown of Hfq was indeed lethal in this strain (44). Conversely, *hfq* mutants are viable in *A. baumannii* ATCC17978, a less virulent and not naturally competent strain, albeit with a profound fitness cost (50, 51). In any case, Hfq plays an important role for *A. baumannii* physiology and virulence, suggesting that small RNAs are key regulators in these processes (33, 48, 50, 52, 53).

While the importance of Arp in controlling natural transformation in *A. baumannii* has been established in this study, the questions about the biological role of Arp *in vivo* remains. What environmental cues are critical for the induction or repression of competent *A. baumannii* cells and what is the effect on cell and population fitness? Multiple signals that trigger natural competence have been uncovered in Gram-negative species, including host-associated signals, carbon source limitation, specific ions, quorum sensing signals, and DNA-damaging agents (14) however, whether any of those are key for Arp-mediated regulation of natural transformation *A. baumannii* remains to be identified. Several studies suggested that competence within this organism is influenced by osmolarity (20, 54, 55). Additionally, a microarray-based study revealed that iron-limiting conditions downregulate multiple T4P genes and reduce swarming motility. However, we did not observe elevated levels of Arp-*pilA* mRNA chimeras during low-iron shock conditions (33, 56). The artificial sweeteners saccharin and acesulfame-K were also shown to repress T4P which impaired twitching motility and DNA uptake in *A. baumannii* AB5075 (57, 58). Future work could focus on the role of Arp in modulating *A. baumannii* competence in response to hyperosmotic stressors or during iron starvation to establish the biological importance of Arp.

A recent ChIP-seq analysis in *A. baumannii* found that the transcription factor BfmR of the two-component system BfmRS may be a direct activator of Arp transcription (38). Additionally, this study found that BfmR directly represses the expression of the PilMNOP complex, proteins that comprise vital components of the *A. baumannii* T4P. The authors did not find that BfmR represses transcription of *pilA*, suggesting that it may do so indirectly, by upregulating expression of Arp as an essential component of the “non-coding arm” of the BfmR regulon. The BfmRS two-component regulatory system is integral in the response of *A. baumannii* to environmental stressors, for instance desiccation, antibiotic shock and survival within hosts, along with motility (59–63). Future work should expand on the integration of Arp within the BfmRS regulatory architecture.

Taken together, this study expands the known repertoire of bacterial sRNAs that regulate horizontal gene transfer and highlights Arp as a promising model for understanding post-transcriptional repression of DNA uptake in *Acinetobacter*. Given the role of natural transformation in the spread of antibiotic resistance, these insights may inform future strategies to disrupt gene acquisition in clinical settings. As Arp represses DNA uptake in a sequence-specific manner, it may serve as a drug target, e.g. for antisense oligonucleotides, to artificially suppress competence in *A. baumannii* (64, 65). This could open novel therapeutic avenues to reduce horizontal gene transfer and curb the spread of antimicrobial resistance in this high-priority pathogen.

## Material and Methods

### Bacterial strains and growth conditions

*Acinetobacter baumannii* AB5075 and *Escherichia coli* TOP10 (Invitrogen) were cultured in lysogeny broth (Lennox, L-broth, 10 g/l tryptone, 5 g/l yeast extract, 5 g/l NaCl) (66). Single, opaque colonies were inoculated in L-broth overnight (16 h) in test tubes prior to inoculating larger culture volumes. Media were supplemented with tetracycline (12 μg/ml), ampicillin (150 μg/ml), chloramphenicol (25 μg/ml), apramycin (60 μg/ml) and sucrose (20% w/v) as required. Motility medium (0.5% (w/v) Sigma (A9539-500G) agarose, 5 g/l tryptone) was used to assess the impact of Arp expression on modulating twitching motility. Transformation medium (2% (w/v) Euromedex agarose (D3-D), 5 g/l tryptone) was used to analyse the impact of Arp in regulating DNA uptake.

### Generation of *A. baumannii* mutants

The *A. baumannii* AB5075 *arp* (sRNA40) locus (positions 1,763,903 to 1,764,077) was removed using a homologous recombination-based strategy (46). The primers used in this study are outlined in **Table S1**. Briefly, ∼2,000 bp regions up-and downstream of *arp* were amplified by PCR using *A. baumannii* AB5075 genomic DNA as the template. Additionally, the *sacB-aacC4* genes were amplified from the pMHL2 plasmid and a chimeric PCR product composed of *sacB-aacC4* flanked by the upstream and downstream regions was constructed using overlap extension PCR. Wildtype *A. baumannii* AB5075 was transformed with this chimeric PCR product using natural transformation as previously described, enabling selection of apramycin resistant colonies (45, 46). The resulting colonies (*A. baumannii* Δ*arp*::*sacB_aacC4* insertion mutants) were also screened for sucrose sensitivity on L-agar plates without NaCl and 20% (w/v) sucrose and the integration of *sacB-aacC4* into the *arp* locus was confirmed by colony PCR. Following this, a second chimeric PCR product composed of only ∼2,000 bp up- and downstream regions of the *arp* locus was created by overlap extension PCR. *A. baumannii* Δ*arp*:*:sacB_aacC4* was transformed with second chimeric PCR product to generate a scarless *A. baumannii* AB5075 Δ*arp* mutant strain. Colonies were screen to validate sucrose resistance and apramycin sensitivity. The loss of the *arp* locus was validated by PCR. A similar strategy was used to create a mutant *A. baumannii* AB5075 strain (Δ*pilA*:*:sacB_aacC4*, referred to as Δ*pilA* in text). Chimeric PCRs were also used to a construct *pilA*::3×FLAG C-terminal chromosomal fusion as previously outlined (33) was validated by PCR and Sanger sequencing.

### Plasmid constructions

PCR products were routinely purified using EasyPure® PCR kit (TransGen Biotech) and plasmids with EasyPure Plasmid MiniPrep Kit (TransGen Biotech) according to the manufacturer’s manual. Recombinant plasmids used in this study, including the pP_L_, pXG10sf, pAMCK14, pAMCK18 plasmids, were constructed using the SLiCE cloning procedure (39, 40, 42, 67). As previously outlined, all DNA oligonucleotides used to amplify insert regions were modified at 5′- and 3′-ends to include ∼20 bp of homology to the ends of the linearised plasmid backbones that were generated by PCR (33). The amplified linear plasmid backbone PCR products were *Dpn*I-digested with Anza™ 10 *Dpn*I (Invitrogen) to remove residual template plasmid DNA prior to performing SLiCE reactions. The pP_L_-Arp plasmid was prepared by amplifying the AB5075 *arp* locus (166 nt), from the predicted TSS identified by differential RNA-seq to a region downstream of the predicted transcriptional terminator (34). This insert was cloned into the linearised pP_L_ plasmid backbone using SLiCE. The pP_L_-Arp* plasmid was created by amplifying up- and downstream regions (∼300 nt) of the Arp interacting region with primers designed to insert point mutations within the predicted interacting sequence. An overlap extension PCR was then used to create a chimeric PCR product composed of both regions. This chimeric PCR product was used as a template for amplifying the pP_L_-Arp* insert and as a template for *in vitro* transcription of Arp* (see below). These same insert PCR products were used to create pAMCK14-Arp and pAMCK14-Arp*. The pXG10-PilA-sfGFP plasmids was prepared by amplifying the *pilA* locus (102 nt) including the TSS and the predicted interaction site. The PilA-sfGFP translational fusion included the 5′-UTR and first 17 codons of *pilA* mRNA translationally fused to sfGFP. This insert was then cloned into the linearised pXG10sf plasmid backbone using SLiCE. A compensatory mutation in *pilA* (*pilA*\*) was created in a similar manner to Arp*. This was used as a template to construct pXG10-PilA*-sfGFP and for *in vitro* transcription of *pilA*\* (see below) and to create pAMCK18-PilA.

### RNA isolation and northern blotting

To assess Arp expression in different conditions and growth stages, RNA from *A. baumannii* was isolated from cells grown to mid exponential phase (MEP, OD_600nm_ = 0.3), late exponential phase (LEP, OD_600nm_ = 1), early stationary phase (ESP, OD_600nm_ = 2) and late stationary phase (LSP, 16 h growth) in L-broth under standard conditions (220 rpm, 25 ml in 250 ml Erlenmeyer flasks, 37°C). Total RNA for use in northern blotting was isolated using TRIzol as previously described (33). Arp expression was analysed using northern blotting as previously described (68). Briefly, total RNA (5 µg) was separated on a 6% (vol vol^−1^) PAA-7 M urea gel at 300 V for ∼2 h, electro-blotted onto an Hybond-N+ membrane (GE Healthcare Life Sciences) and then UV-crosslinked. Membranes were pre-hybridized in Hybri-Quick buffer (Carl Roth AG) and incubated with ^32^P-labeled probes at 42 °C. Blots were washed three times with SSC buffer (5×, 1×, 0.5×), exposed and then visualized on a phosphorimager (FLA-3000 Series, Fuji).

### *In vitro* transcription and RNA radiolabelling

Radiolabelled RNA was generated by *in vitro* transcription as described previously (68). Templates were PCR-amplified from genomic DNA using T7 promoter-encoding primers. To produce the Arp* mutant variant, a modified template was prepared by overlap extension PCR introducing the desired mutations (substitution of CCCC→GGGG, at positions 51–54). *In vitro* transcription was performed using the MEGAscript T7 Kit (Invitrogen), followed by DNase I treatment (1 U, 37°C, 15 min) to remove template DNA. Transcription products were separated on a 6% polyacrylamide gel (7 M urea, 1× TBE), alongside a LowRange RNA ladder (ThermoFisher Scientific). The relevant RNA bands were excised, and RNA was eluted overnight at 8°C with shaking (1400 rpm) in elution buffer (0.1 M NaOAc, 0.1% SDS, 10 mM EDTA). Eluted RNA was recovered by ethanol precipitation (ethanol:NaOAc, 30:1), washed with 75% ethanol, and resuspended in DEPC-treated water. For 5′ end-labelling, 50 pmol of RNA was dephosphorylated with calf intestine alkaline phosphatase (25 U, 37°C, 1 h; Invitrogen) and extracted with phenol:chloroform:isoamyl alcohol (25:24:1).

### Electrophoretic mobility shift assay (EMSA)

RNA-RNA EMSAs were done as previously described (68). Briefly, reactions were prepared containing 1x RNA structure buffer (SB; Ambion), 1 µg yeast RNA (∼4 µM), and 5’ end-labelled RNA (4 nM). The in-vitro-transcribed target mRNA was supplied at 0; 8; 16; 32; 64; 128; 256; 512; and 1,024 nM. Reactions were incubated for 1 h at 37 °C and stopped with 3 µL of 5x native loading dye (0.2% bromophenol blue, 0.5x TBE, 50% glycerol) and ran on a native 6% (vol vol−1) PAA gel in 0.5x TBE buffer at 4 °C at 300 V for 3 h. The gel was dried (Gel Dryer 583, Bio-Rad) exposed and visualized on a phosphorimager (FLA-3000 Series, Fuji).

### In-line probing

Assays and the preparation of sequencing ladders were done as previously described (68), using 0.4 pmol labelled Arp RNA. In-line probing reactions were incubated for 40 h at room temperature in 1x in-line probing buffer (100 mM KCl, 20 mM MgCl2, 50 mM Tris-HCl, pH 8.3). Where required, target *pilA* RNA was supplied at 0.2, 0.4 and 2 pmol. All reactions were stopped with colourless loading dye (1.5 mM EDTA, pH 8, 10 M urea). Then samples were run on a 10% (vol vol−1) PAA-7 M urea sequencing gel and visualized as described above.

### Confirmation of sRNA-mRNA interactions in *E. coli*

Plasmids constitutively expressing Arp (pP_L_-Arp) or Arp* (pP_L_-Arp*) were constructed and the impact of Arp/Arp* overexpression on PilA-sfGFP (pXG10sf-pilA) was assessed by measuring PilA-sfGFP fluorescence intensity compared to a control strain co-expressing a control plasmid (pJV300). Individual *E. coli* colonies were used to inoculate 5 ml L-broth containing ampicillin and chloramphenicol. Overnight cultures (1:1000 dilutions) were inoculated into 25 ml flasks of L-broth containing ampicillin and chloramphenicol. The next day, cultures were diluted 1:1000 into fresh 25 ml L-broth containing ampicillin and chloramphenicol in a 250 ml Erlenmeyer flask and incubated at 37°C and 220 rpm agitation for 16 h. The cultures were then aliquoted into 96-well plates and fluorescence (arbitrary units) and optical density (OD_600_) were measured in a Synergy H1™ Hybrid Multi-Mode microplate fluorometer (Bio-Tek) with excitation (485 nm) and emission (508 nm). The fluorescence of strains was compared by quantifying the observed fluorescence relative to the optical density at 600 nm.

### Confirmation of sRNA-mRNA interactions in *A. baumannii*

Plasmids constitutively expressing Arp (pAMCK14-Arp) or Arp* (pAMCK14-Arp*) were constructed and the impact of Arp/Arp* overexpression on PilA-sfGFP (pAMCK18-pilA) was assessed by measuring PilA-sfGFP fluorescence intensity compared to a control strain co-expressing a control plasmid (pAMCK15). Individual *A. baumannii* Δ*arp* colonies were used to inoculate 4 ml L-broth containing apramycin and tetracycline. These cultures were grown overnight and 5 μl of these cultures were inoculated into 1 ml fresh L-broth (containing apramycin and tetracycline) and incubated at 37°C and 220 rpm agitation for 30 h. The cultures were then aliquoted into 96-well plates (company) and fluorescence (arbitrary units), and optical density (OD_600_) were measured in a Synergy HTX Hybrid Multi-Mode microplate fluorometer (Bio-Tek) with excitation (460/40 nm) and emission (528/20 nm). The fluorescence of strains was compared by quantifying the observed fluorescence relative to the optical density at 600 nm.

### Sequence conservation analysis

A phylogenetic analysis was performed to assess the conservation of the *arp* sequence across the *Acinetobacter* genus. A BLASTn search (v2.12.0+) of the *A. baumannii* AB5075 *arp* sequence was conducted against 923 complete *Acinetobacter* genomes available as of April 2024, using default parameters (69). Homologous sequences showing >95% identity and >90% alignment length were retained for downstream analysis. Multiple sequence alignment was performed using MUSCLE (v5.2, linux64) with default settings (70). A maximum likelihood phylogeny was inferred using IQ-TREE multicore version 2.0.7 with Extended Model Selection (-m MFP) and 2,000 ultrafast bootstrap replicates (71). The resulting tree was visualized using iTOL (72).

### Protein isolation and immunoblotting

Western blotting experiments were used to compare PilA::3×FLAG expression levels in wildtype and Δ*arp* strains. Individual colonies of these strains were used to inoculated 1 ml of motility medium. These motility medium tubes were then incubated at 37°C for 24 h before cells were resuspended in 200 µl PBS. These cells were then washed twice in PBS and collected by centrifugation (5 min, 2,500 g, 4 C) before being lysed in Lämmli buffer at 95°C for 10 min. Equivalents of OD_600_ 0.03 cells were then separated on 15% (w/v) discontinuous SDS-polyacrylamide gels. The separated proteins were then transferred to a nitrocellulose membrane (AmershamTM Protran 0.45 μm, Cytiva), and PilA:::3×FLAG expression was detected by anti-FLAG antibodies (1:10,000, Sigma #F3165) while DnaK loading control was detected by anti-DnaK antibodies (1:5,000, Enzo #8E2/2). Both primary antibodies were detected using rabbit anti-mouse IgG, 94 HRP-conjugate (1:10,000, Merck #AP160P). An ImageQuant LAS4000 imager (GE Healthcare) performed chemiluminescent imaging. Quantification was performed using ImageJ software (73).

### Twitching motility assays

Twitching motility assays were performed using a protocol that was adapted from previous studies (74). Briefly, individual colonies of similar sizes were stabbed through the centre of motility medium containing petri dishes. The plates were then placed in sealed bags and incubated upright at 37°C for 5 days. The zones of motility were subsequently quantified using ImageJ imaging software (73).

### DNA uptake assays

Natural transformation efficiency was evaluated by measuring the ability of *A. baumannii* strains to acquire and maintain the pAMCK18 plasmid, following previously established protocols (45, 46). Briefly, single colonies were inoculated into 2 mL of L-broth supplemented with apramycin and grown at 37°C for 3 hours. Cultures were then diluted to an OD₆₀₀ of 0.01 to standardize cell densities. For transformation, 5 µL of the diluted culture was mixed with 20 ng of plasmid DNA, and 2.5 µL of this mixture was spotted onto transformation medium. Following overnight incubation at 37°C, cells were resuspended in 100 µL of PBS, serially diluted, and plated onto L-agar with or without tetracycline and apramycin. Transformation efficiency was calculated as the proportion of tetracycline-resistant colonies relative to the total viable population.

## Author contributions

FJH and CK designed the study. FJH and CK wrote the original draft which all other authors edited. FJH, TG, OC carried out lab experiments. KH created the Arp browser. ASE carried out bioinformatic analyses. FJH, TG, MND, ASE, ABF, MJG, AJW and CK analysed the data.

## Acknowledgments and funding sources

F. Hamrock was supported by a 1252 Postgraduate Research Studentship from Trinity College Dublin, a 2023 Irish Research Council, Government of Ireland Postgraduate Scholarship (GOIPG/2023/4991) and was a recipient of an EMBO Scientific Exchange Grant (no. 9316). M. Gebhardt was supported by NIH grant GM156848. Deirdre Muldowney (TCD) is acknowledged for technical support.

