## Supplementary Data for "DNA uptake and twitching motility are controlled by the small RNA Arp through repression of pilin translation in *Acinetobacter baumannii*"

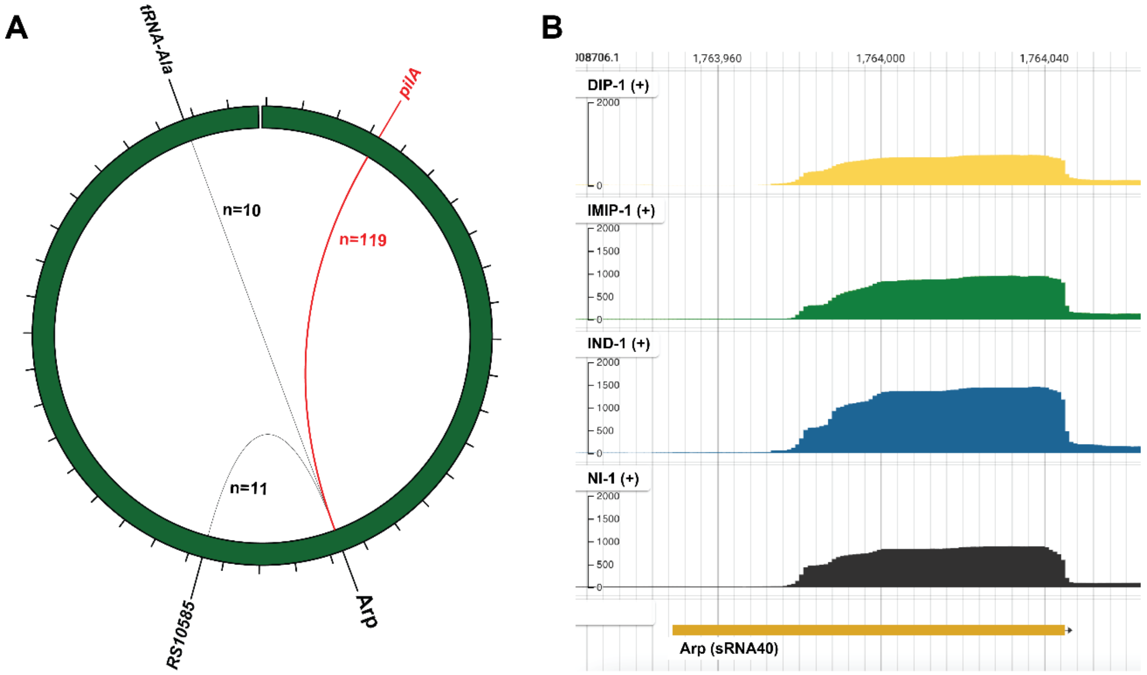


**Supplementary Figure 1 – Arp-*pilA* interactions are prevalent in Hi-GRIL-seq.** (**A**) Hi-GRIL-seq (1) revealed that the *A. baumannii* sRNA Arp formed abundant chimeras with *pilA* mRNA (n=119 chimeras). Arp interactions are shown across the *A. baumannii* AB5075 chromosome where the line thickness is proportional to the depicted number of chimeric reads obtained. (**B**) Visualisation of the of the *arp* locus read coverage in the Hi-GRIL-seq mapped (non-chimeric) reads using JBrowse2 (2). One biological replicate of each Hi-GRIL-seq condition used is shown ((NI = non-induced control, IND = induced sample, DIP = iron starvation and IMIP = imipenem shock).

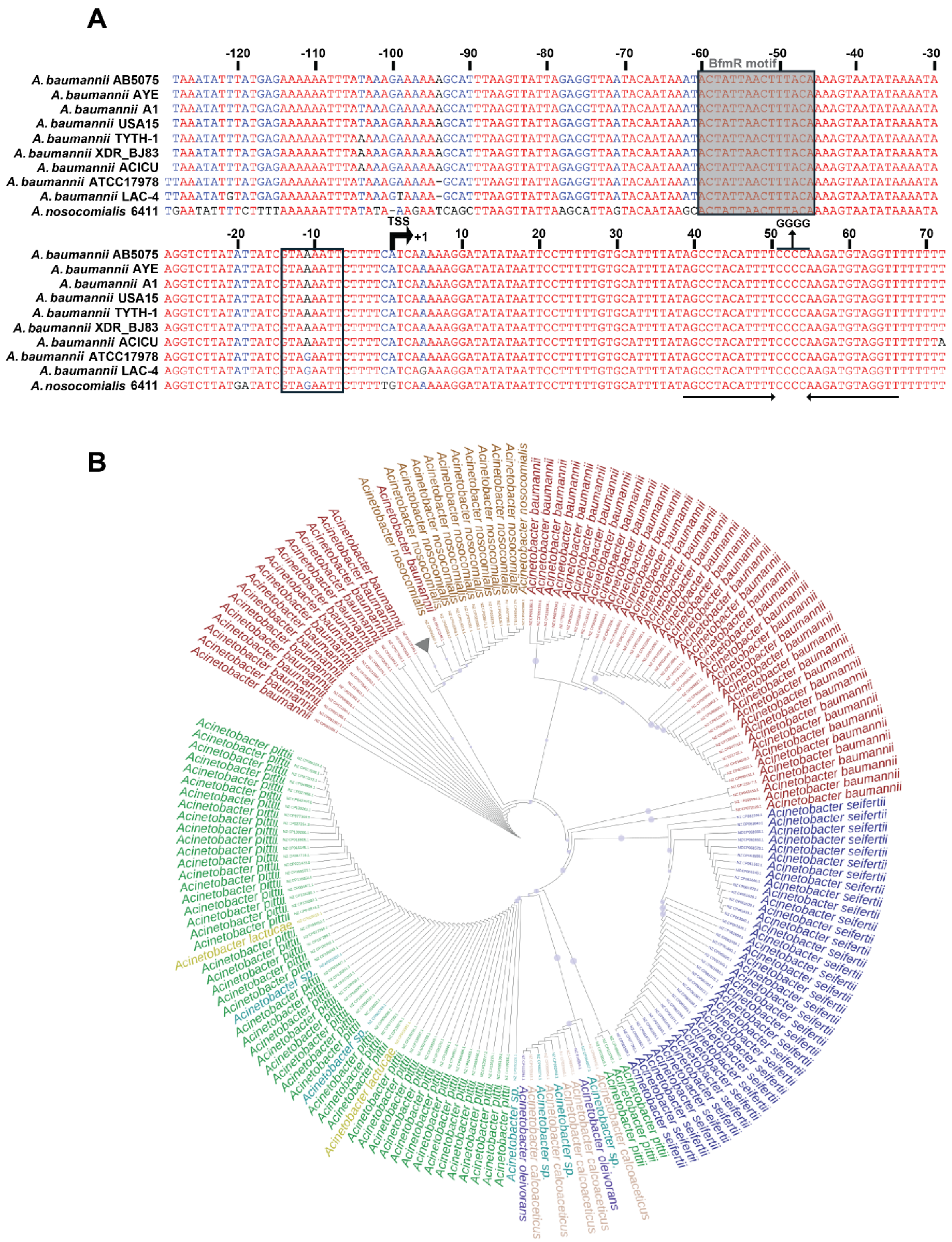


**Supplementary Figure 2 – Arp is broadly conserved among *Acinetobacter* species.** (A) Sequence alignment of *arp* across diverse *Acinetobacter* species. Alignment was performed using the Multalin tool (3), with nucleotide coloration indicating conservation levels. The predicted transcriptional start site (TSS) is denoted as +1, with nucleotide positions shown relative to this site. The predicted interacting region is underlined, and its mutated version is also indicated. Putative −10 promoter element is boxed, and the predicted Rho-independent terminator is marked by paired arrows below the sequence. The putative BfmR binding motif previously identified is boxed and is shaded in grey (4). (**B**) Phylogenetic analysis of *arp* homologues across the *Acinetobacter* genus. A maximum likelihood phylogenetic tree was constructed from 725 *arp* homologues identified in complete *Acinetobacter* genomes. Sequences with >95% identity and >90% alignment length to the *A. baumannii* AB5075 query were included. Bootstrap support values are indicated by circles at branch nodes; no branches exceeded 90% support, consistent with limited sequence divergence. For clarity, a clade containing 551 highly similar *A. baumannii* and *Acinetobacter sp.* sequences was collapsed (grey triangle).

**A**

| Motif | Sequence | Position within Arp (nts) |
|---|---|---|
| ARN1 | AAA | 4 – 6 |
| ARN2 | AAA | 5 – 7 |
| ARN3 | AAA | 6 – 8 |
| ARN4 | AAG | 7 – 9 |
| ARN5 | AGG | 8 – 10 |
| ARN6 | AAU | 17 – 19 |
| ARN7 | AGC | 39 – 41 |
| ARN8 | AAG | 55 – 57 |
| ARN9 | AGA | 56 – 58 |
| ARN10 | AGG | 62 – 64 |

**B**

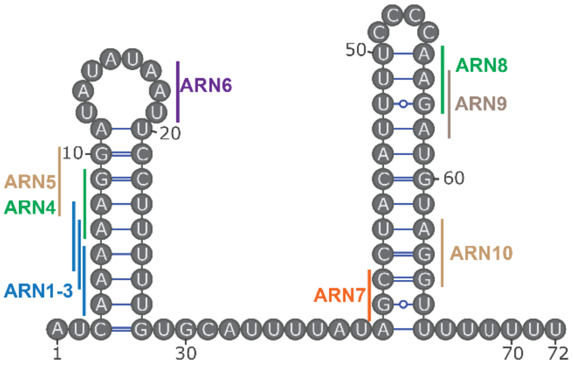


**Supplementary Figure 3 – Identification of ARN motifs in Arp.** (**A**) Table listing the ARN motifs (consensus: A-A/G-N) identified in the Arp sRNA. (B) Secondary structure representation of the Arp sRNA with annotated locations of ARN motifs (ARN1–10). Motifs are marked according to their position in the primary sequence and clustered where overlapping. Each motif is colour-coded based on its exact nucleotide sequence.

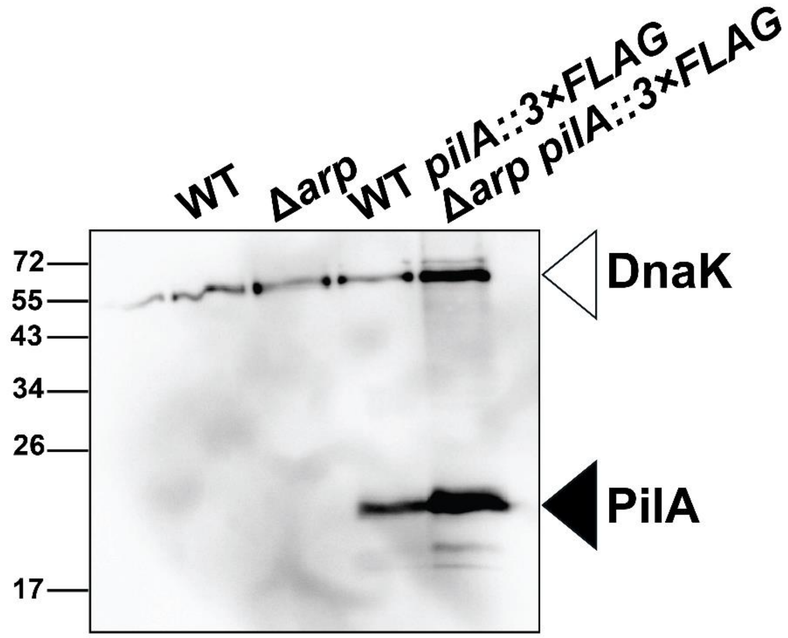


**Supplementary Figure 4 – Validation of chromosomally FLAG-tagged PilA expression in *A. baumannii*.** Whole-cell lysates were prepared from wildtype (WT) and Δ*arp A. baumannii* strains carrying a chromosomal *pilA::3×FLAG* fusion and grown to mid-exponential phase. WT and Δ*arp* strains lacking the FLAG fusion were included as negative controls. PilA::3×FLAG was detected by western blotting using anti-FLAG antibody. DnaK was probed as a loading control.

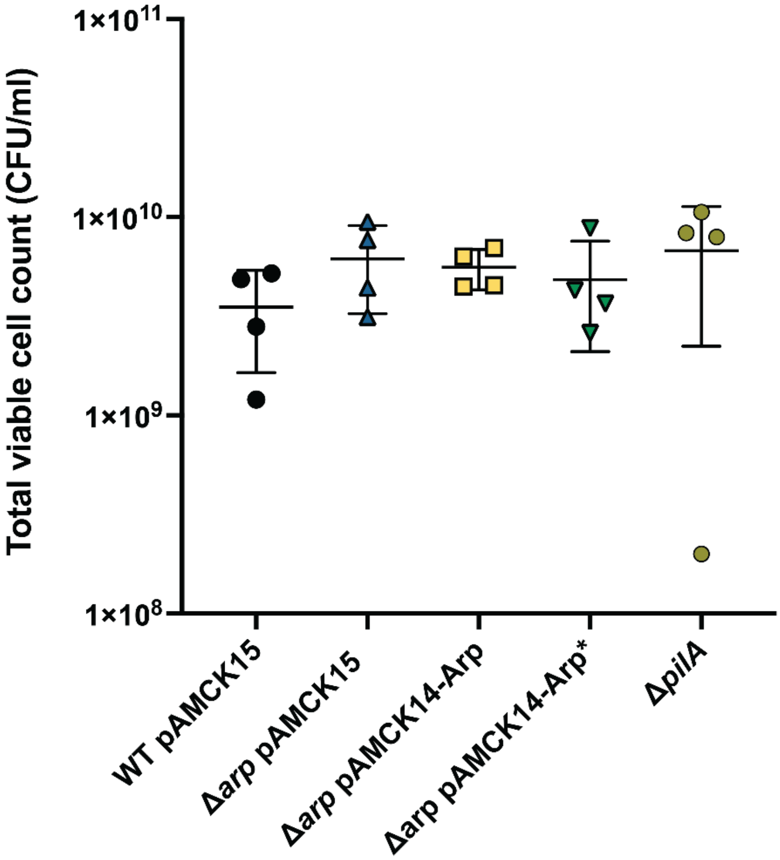


**Supplementary Figure 5 – Natural transformation controls.** The total viable cell counts recovered from the natural transformation assay on non-selective L-agar. *A. baumannii* AB5075 wildtype (WT) and Arp-deletion strains (Δ*arp*) carrying either an empty vector (pAMCK15), an Arp overexpression plasmid (pAMCK14-Arp), or a interacting region mutant of Arp (pAMCK14-Arp*). A Δ*pilA* strain was included as a negative control. Error bars represent the standard deviation of four independent biological replicates (n=4). Statistical analysis was performed using one-way ANOVA. No statistically significant differences were observed.


1. F. J. Hamrock, *et al.*, Global analysis of the RNA–RNA interactome in Acinetobacter baumannii AB5075 uncovers a small regulatory RNA repressing the virulence-related outer membrane protein CarO. *Nucleic Acids Res.* **52**, 11283–11300 (2024).
2. C. Diesh, *et al.*, JBrowse 2: a modular genome browser with views of synteny and structural variation. *Genome Biol.* **24**, 74 (2023).
3. F. Corpet, Multiple sequence alignment with hierarchical clustering. *Nucleic Acids Res.* **16**, 10881–10890 (1988).

4. N. Raustad, *et al.*, A phosphorylation signal activates genome-wide transcriptional control by BfmR, the global regulator of Acinetobacter resistance and virulence. *Nucleic Acids Res.* **53**, gkaf063 (2025).
